# Antibody-mediated antigen tethering as a regulator of germinal-center B cell affinity and diversity

**DOI:** 10.64898/2026.01.14.699581

**Authors:** Colin Scheibner, Andrew G.T. Pyo, Dennis Schaefer-Babajew, Ned S. Wingreen

**Affiliations:** Joseph Henry Laboratories of Physics, Princeton University, Princeton, New Jersey 08544, USA; Princeton Center for Theoretical Science, Princeton University, Princeton, New Jersey 08544, USA; Department of Applied Physics, Stanford University, Stanford, CA 94305, USA; Laboratory of Molecular Immunology, Rockefeller University, New York, NY 10065, USA; Department of Internal Medicine, Yale School of Medicine, New Haven, CT 06520, USA; Lewis-Sigler Institute for Integrative Genomics, Princeton University, Princeton, New Jersey 08544, USA; Department of Molecular Biology, Princeton University, Princeton, New Jersey 08544, USA; Weill Cancer Hub East Investigator

## Abstract

To optimally protect a host, the adaptive immune system must generate antibodies (Abs) that bind the pathogen with high affinity, yet are diverse enough to protect against potential escape mutations. How the immune system can steer between affinity and diversity remains an open question. During the immune response, B cells inside germinal centers (GCs) compete for survival based on their ability to extract antigen from the surface of specialized presenting cells called follicular dendritic cells (FDCs). Curiously, FDCs tether antigen via not one, but two qualitatively distinct routes: direct tethering, in which the linking protein is covalently bound to the Ag, and indirect, Ab-mediated tethering, in which the linking protein is an Ab itself. Using agent-based simulations motivated by the biophysics of Ag extraction, we suggest that the dual-tethering system can improve the robustness of affinity maturation, and may provide a mechanism to steer the balance between affinity and diversity of the GC B cell population. Our model is consistent with experimental observations reporting low-affinity but high-diversity GCs in the absence of Fc*γ*R, a key Ab-tethering receptor on FDCs.

---

From the dazzling array of cell types to the diversity of receptors and signals, how the molecular complexity of the adaptive immune system gives rise to robust immune function remains a conceptual challenge. In particular, a vital feature of the adaptive immune system is its ability to generate novel, protective antibodies (Abs) when faced with a specific threat. Mechanistically, this process— affinity maturation—occurs in microanatomical environments known as germinal centers (GCs) [1–3] (Fig. 1a). Briefly, GCs are organized into two compartments: the light zone (LZ) and the dark zone (DZ). In the LZ, antigen (Ag) is presented on the surface of follicular dendritic cells (FDCs) [4–6]. B cells explore the FDCs for cognate Ag, and upon detection, a B cell attempts to extract the Ag using its B cell receptor (BCR) [7–15]. If extracted, the Ag is endocytosed, digested into peptides, and presented on MHC class II molecules to follicular helper T cells. If sufficient T-cell help is received, the B cell migrates to the DZ, where it undergoes rapid divisions and somatic hypermutation [16, 17]. The number of divisions is set by the amount of T-cell help [18]. The mutated daughters then return to the LZ and repeat this process, by default undergoing apoptosis if they fail either to find cognate Ag or to receive T-cell help. Hence, B cells that are able to acquire more Ag and therefore receive more T-cell help will produce more progeny and thus form a larger fraction of the future GC population. During this process, a small fraction of B cells are exported to effector fates, including plasma cells (responsible for immediately secreting Abs) and memory cells (poised for reactivation during future infections).

**FIG. 1.**
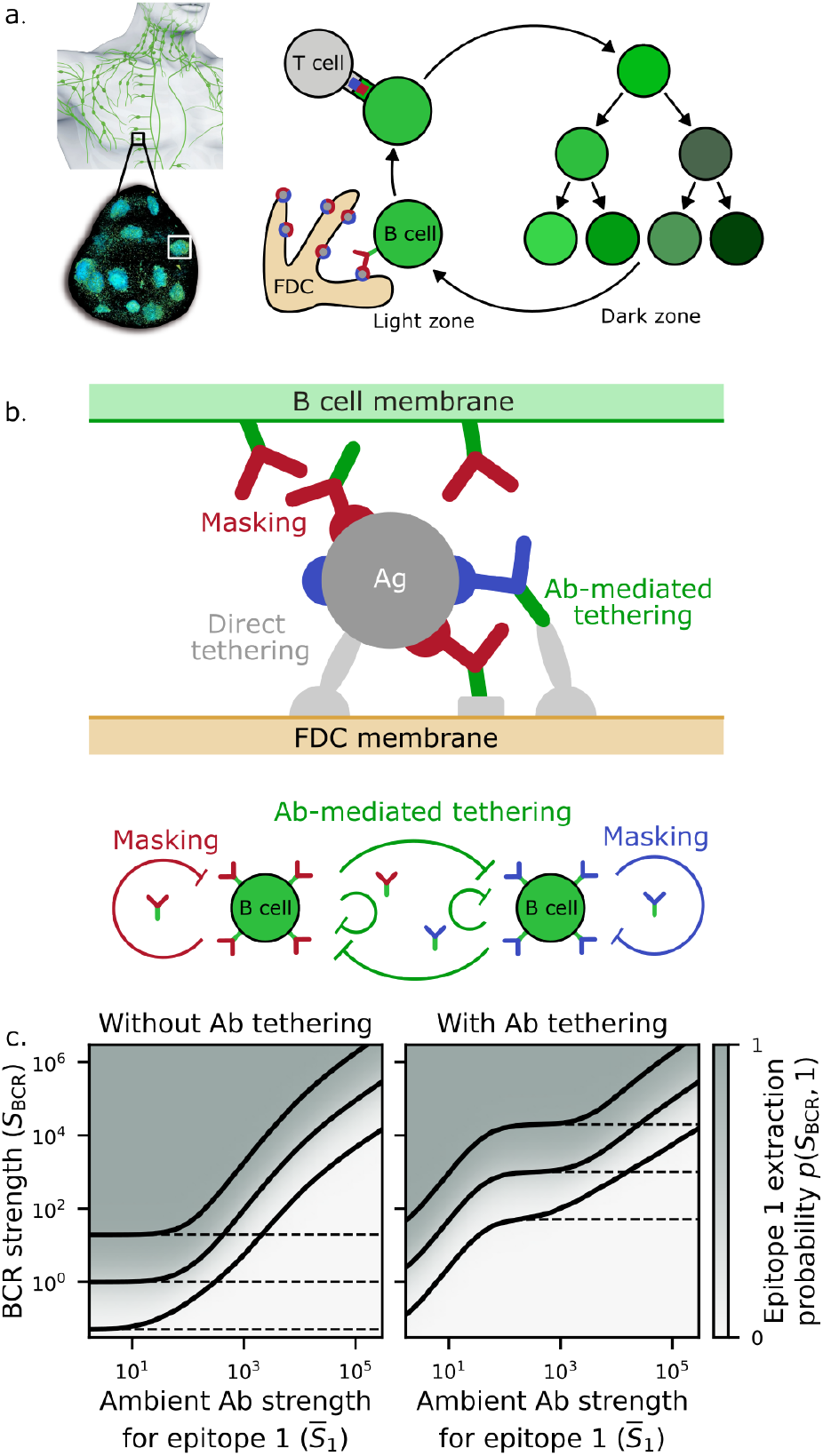
The germinal center (GC) reaction and mechanisms of Ag tethering. **(a)** Left: schematic of the human lymphatic system and light-sheet fluorescence micrograph of a lymph node, with individual GCs marked by fluorescent labeling of proliferating B cells (adapted from [53]). Right: the GC reaction. A B cell extracts Ag from an FDC via its BCR, presents pMHC to a follicular helper T cell, and then migrates to the DZ for proliferation and somatic hypermutation (color change). **(b)** Upper: schematic of the B cell–FDC synapse. The IC, containing Ag with two epitopes (red and blue) coated in cognate Abs, is tethered to an FDC either directly through complement fragments (gray ellipse) bound to complement receptors (CRs; semicircles), or indirectly through Ab-mediated interactions involving Fc*γ*Rs (gray rectangle) or complement-bound Abs. Soluble Abs can also mask epitopes and inhibit BCR binding. Lower: conceptual schematic. Epitope masking reduces Ag acquisition for B cells of the same epitope class; Ab-mediated tethering reduces Ag acquisition for all B cells. **(c)** The Ab feedback model (Eq. (1)) with and without Ab tethering. Heatmaps: probability that a B cell targeting epitope 1 extracts a unit of Ag as a function of BCR strength (*S*_BCR_) and soluble Ab strength against epitope 1 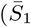; Eq. (2)), assuming 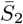 = 0 for illustration. Solid black contours denote 5%, 50%, and 95% extraction probability for epitope 1. Dashed contours show the corresponding probabilities for epitope 2. Notably, Ab tethering (right) influences extraction probability for both epitopes. Parameters: *S*_dir_ = 1; *S*_unbound_ = 10^4^; *S*_indir_ = 0.2; *A*_indir_ = 0 (left), 5 *×* 10^3^ (right). (Schematic of lymphatic system in (a) adapted from Stocktreck Images.)

To optimally protect the host, the GC reaction must achieve two competing objectives: the evolved Abs must bind with high enough affinity to clear the pathogen, but the Ab repertoire must also be diverse enough to anticipate future escape mutations [19]. A key factor shaping this balance is “Ab feedback”: the influence of previously secreted Abs on ongoing GC selection [20–36]. Briefly, Ag is generally presented on the membrane of an FDC in the form of an immune complex (IC), consisting of one or more Ags coated in complement (proteins of the innate immune system) and Abs (see Fig. 1b). As the immune response progresses, weaker Abs in ICs are displaced by stronger Abs. This turnover has been proposed, and in some cases demonstrated, to influence the dynamics of the GC through multiple mechanisms. For instance, upon repeated exposure (e.g., sequential vaccinations), the total amount of Ag presented in the GC increases on later doses because more Abs pre-exist to sequester and preserve the Ag [37]. Such mechanistic insights have led to proposals for optimal vaccine dosing [37–43], and have helped explain why three doses of the COVID vaccine provide broader protection than two [44].

Beyond increasing the amount of presented Ag, Ab feedback is also thought to affect the relative selection pressure during a GC reaction. Different lineages of B cells can bind to different sites, known as epitopes, on the Ags. IC Abs can block BCRs that compete for the same or partially overlapping epitopes. This “epitope masking” has alternatively been proposed to steer GC selection in several distinct ways [20–22, 24, 26, 28, 33, 35, 44]. When only a single epitope class is present, masking is thought to drive the GC B cells to higher affinity by incrementally increasing the difficulty for BCRs to bind to Ag [20]. With multiple epitopes, epitope masking has instead been argued to drive diversification of GCs (by giving a competitive edge to B cells cognate to alternative epitopes [22, 24, 28, 44]). Whether the immune system has evolved a robust mechanism to steer between these two fates—increasing affinity or diversification—remains an open question.

In addition to masking, Ab feedback is also thought to affect the strength with which ICs are bound to FDCs. Ag is tethered to FDCs through two qualitatively distinct mechanisms. The most widely recognized is direct tethering, in which a complement protein (namely C3d) covalently binds to Ag and binds to complement receptors (CRs) on FDCs [4, 45, 46]. The second is Ab-mediated tethering, in which the IC is noncovalently bound by a soluble Ab that is itself connected to the FDC membrane—either through an Fc*γ*R (specifically Fc*γ*RIIb) [47–51], which binds to the conserved tail of IgG Abs, or through complement fragments covalently bound to the Ab [52]. Jiang and Wang proposed a conceptual model in which only Ab-mediated tethering is present [27]. In this picture, Ab feedback drives GC selection to higher affinity by gradually increasing the BCR affinity required to extract ICs from the FDCs. Why the immune system nevertheless evolved to use two distinct tethering mechanisms, and how this interacts with masking, has not been addressed.

Here, we theoretically investigate GC selection in light of all three effects: masking, direct tethering, and indirect, Ab-mediated tethering. Our results suggest that all three mechanisms may collaborate to provide robust affinity maturation. Without Ab-mediated tethering, for instance, we find that epitope masking alone leaves the GCs susceptible to invasion by B cells that target sub-dominant epitopes with low affinity, and therefore are not yet self-inhibited by epitope masking. By contrast, Ab-mediated tethering increases the difficulty of Ag extraction for all B cells, stabilizing affinity maturation for B cells targeting a dominant epitope with high affinity. While further experimental investigation is needed, these results suggest that indirect, Ab-mediated tethering could be used by FDCs as a biophysical control knob to actively regulate the affinity–diversity tradeoff.

## Model

We investigate the interplay between epitope masking, direct Ag tethering, and Ab-mediated Ag tethering using a mathematical model of GC selection. As detailed in the Methods, the model consists of agent-based GC selection with proliferation and mutations coupled to a phenomenological description of Ab feedback. In brief, the GC model comprises a population of B cells (the agents), each in one of three compartments, the LZ, DZ, or plasma B cell compartments. In the LZ, each B cell attempts to extract Ag at a given rate, and when an attempt is successful, the B cell acquires a unit of Ag. Simultaneously, LZ B cells are stochastically inspected by T cells; upon inspection, those that have collected a sufficient amount of Ag are sent to the DZ to undergo a number of cell divisions determined by their Ag load. During these divisions, the DZ B cells acquire mutations which affect the quality of each daughter’s BCR. The mutated daughters then return to the LZ and the process continues. The LZ B cells that do not present enough Ag to be sent to the DZ by the T cells eventually undergo apoptosis. During this reaction, a small fraction of B cells are exported to the plasma compartment.

While the details of each step of the above reaction scheme continue to be subject to intense experimental and theoretical investigation, we focus on the stability of affinity maturation in relation to epitope masking, direct tethering, and Ab-mediated tethering. We conceptualize each B cell receptor as having a *strength S*_BCR_ for extracting Ag. This dimensionless strength may be thought of as a proxy for thermodynamic affinity (inverse dissociation constant), but the ability to extract Ag in context is also influenced by other biophysical factors (expression levels, molecular flexibility, epitope valency, etc.), so for clarity, we give it a distinct name. When a B cell encounters an FDC, the B cell’s probability of extracting Ag, *p*(*S*_BCR_, *i*), depends on its BCR strength *S*_BCR_, the cognate epitope *i*, and the state of the secreted antibodies. Rather than directly coarse-graining the underlying biophysics of Ag extraction, we consider a phenomenological form of *p*(*S*_BCR_, *i*) which captures the underlying scales and processes.

In particular, for a B cell to have a high probability of extracting an Ag, it must be stronger than two scales: a tethering scale *S*_tether_, which captures how strongly the IC is bound to the FDC, and a masking scale 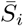 which captures the strength of epitope masking, both of which depend on Ab feedback. A simple functional form realizing this is

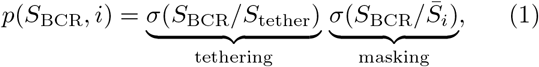

where *σ*(*x*) = *x/*(1 + *x*) is a sigmoid ranging between 0 and 1. Crucially, both 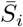 and *S*_tether_ depend on Ab feedback, and they do so in qualitatively distinct ways. For spatially well-separated epitopes, masking primarily inhibits Ag extraction by B cells that target the same epitope as the Ab; by contrast, binding of Ag to the FDC by Ab-mediated tethering makes Ag extraction more difficult for all B cells, regardless of epitope specificity. This inhibition circuit, conceptually summarized in Fig. 1b (lower), is the key ingredient in the following expressions for 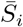 and *S*_tether_.

For epitope *i*, a reasonable choice for the masking scale 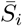 due to ambient antibodies is one that depends only on the exported plasma B cells specific to that epitope and increases with either their typical strength or number. In our model, we take

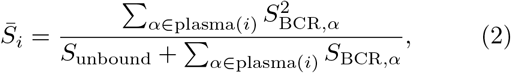

where the sum over *α* plasma(*i*) includes all B cells that have been exported to the plasma-cell fate and target epitope *i*. The quadratic weighting in the numerator is convenient because it is particularly sensitive to the highest-strength plasma cells. If one interprets the strengths roughly as thermodynamic affinities, then Eq. (2) represents an average plasma cell affinity weighted by a likelihood of its secreted Ab being bound. The parameter *S*_unbound_ sets a probability of the epitope being unbound—a nonzero *S*_unbound_ implies that an epitope is not strongly masked when the associated plasma B cells are sufficiently weak or few in number. As either the strength or number of the exported plasma cells increases, the masking by ambient Abs becomes progressively stronger.

Next, a phenomenological expression for the tethering scale *S*_tether_ should include two contributions: direct tethering, which is present even in the absence of Ab feedback, and Ab-mediated tethering, which vanishes when no Abs are present. With these ingredients in mind, we take

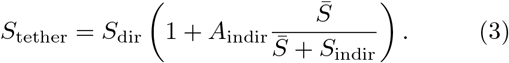

The first factor, *S*_dir_, sets the scale of direct tethering alone, and is naturally independent of Ab feedback. The second is a modulation factor that captures the effect of Ab-mediated tethering. Unlike the masking of a specific epitope *i*, Ab-mediated tethering depends on the total strength 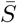 from Abs targeting all epitopes,

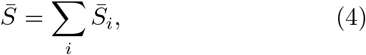

where the sum is over all epitopes *i*. The Ab-mediated tethering term has two phenomenological parameters. First, *S*_indir_ sets the scale at which *S*_tether_ saturates as a function of 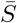. Saturation at some scale is expected because Ab-mediated tethering can only be as strong as the receptors, mechanically in series, that bind the Abs or Ab-bound complement to the FDC membrane. Second, *A*_indir_ sets the magnitude of the overall modulation factor. In the absence of masking, *S*_tether_ can be thought of as an effective barrier to entry in the GC in the sense that a BCR much weaker than *S*_tether_ will not be able to extract Ag. Without Ab-mediated tethering, this barrier to entry resides at the scale *S*_dir_, but as Abs begin to provide additional indirect tethering, the scale is shifted upward, ultimately by a factor of *A*_indir_. Motivated by the fact that the affinity of collaborating bonds tends to multiply rather than add in the fast rebinding limit, direct and Ab-mediated tethering enter as a product in Eq. (3).

To illustrate the difference in *p*(*S*_BCR_, *i*) with and without Ab-mediated tethering, Fig. 1c shows the probability of Ag extraction *p*(*S*_BCR_, 1) as a function of 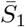 assuming 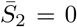. The solid black curves represent 5%, 50%, and 95% probabilities of Ag extraction by a B cell targeting epitope 1. The dashed black curves represent the corresponding probabilities for epitope 2. Without Ab-mediated tethering, B cells targeting epitope 1 face an increasingly difficult Ag-extraction landscape as 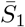 increases due to masking, while epitope-2-specific B cells are unaffected. However, with Ab-mediated tethering, there is a more rapid initial increase in the difficulty to extract Ag as a function of 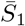, and crucially this initial increase affects B cells of *both* epitope classes.

We emphasize it is the qualitative features of Eqs. (1-4)—rather than their precise functional forms—that are robust to microscopic details. Indeed, Ag extraction is an extraordinarily rich process [7, 8, 10], involving active forces [9, 11–14, 27], membrane mechanics [9], multivalency [54–57], and molecular flexibility and geometry [58–60]. Our purpose here is not to coarse-grain these processes explicitly, but rather to track how the key scales contribute to maturation and/or diversification in the GC reaction. Nonetheless, we note that even a naive first-passage-time approach to extraction (see Supplementary Information) yields the same functional form as Eq. (1-3), which can nevertheless be motivated on phenomenological grounds as done above.

In the following, we explore how this Ag-extraction landscape affects the dynamics and outcomes of the GC reaction. While entrance into a GC is a rich topic [24], in our minimal model, we assume external (“naïve”) B cells continuously arrive in the LZ at a fixed rate. Each new B cell has some probability of targeting either of two epitopes, which we take to be equal for simplicity. While epitopes could differ in many regards (e.g., mutational landscape), for simplicity we consider them differing only in the distribution of precursor B cell strengths. Specifically, we assume an exponential distribution of log-strengths: 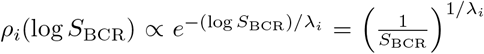, cf. [44], where the decay scale *λ*_*i*_ depends on the epitope (see Methods for additional details). An epitope with larger *λ*_*i*_ is considered immunodominant.

## Results

### Without Ab-mediated tethering, epitope masking promotes epitope diversity and stalls affinity maturation

To illustrate the interplay between epitope masking, direct tethering, and Ab-mediated tethering, we first examine B cell competition when only masking and direct tethering are present (i.e., *A*_indir_ = 0). As a baseline, we first consider a simple situation in which the Ag has only one available epitope class. Figure 2 shows a kymograph of a representative stochastic realization, in which the intensity of the color indicates the number of LZ B cells of strength *S*_BCR_. Also plotted are the masking scale 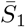 due to the plasma cell output (pink curve) and the *p*(*S*_BCR_, 1) = 5%, 50%, and 95% contours (black curves). Affinity maturation proceeds most effectively when the distribution of B cells roughly overlaps with the 5%-to-95% window; otherwise, the LZ B cells do not experience a reward gradient. For a single epitope subject to masking effects, the 5%-to-95% window shifts upward on the *S*_BCR_ axis as the masking scale 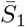 increases due to the plasma cell output. This moving gradient then drives the GC reaction to higher affinity. This scenario is consistent with Ref. [20], which suggests an effective comoving equilibrium between soluble Ab and GC BCR affinities.

**FIG. 2.**
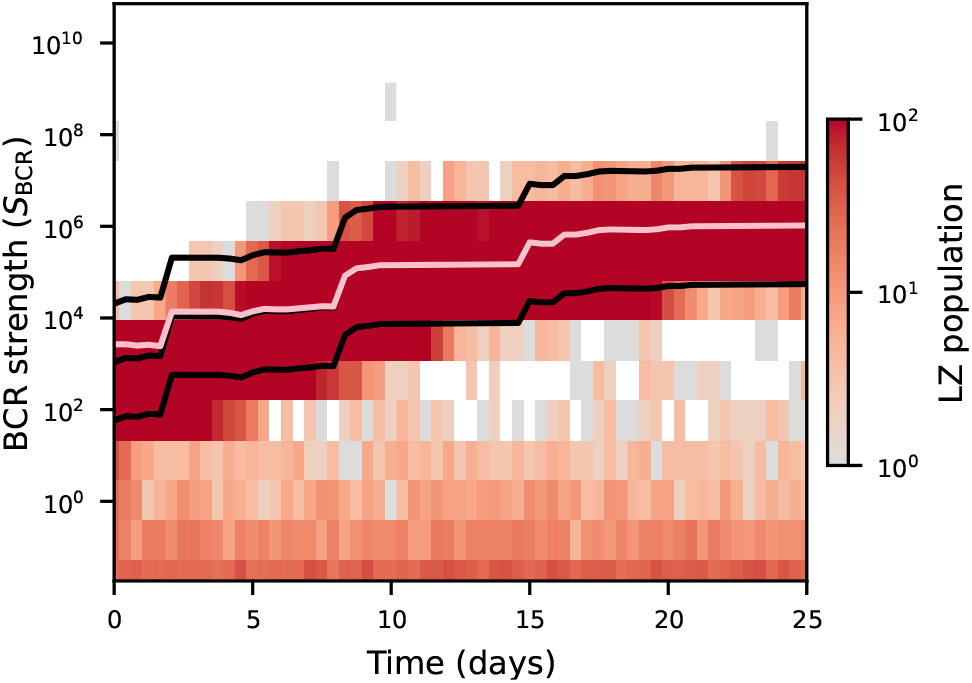
Affinity maturation with a single epitope. Example kymograph from a GC simulation with only a single epitope and no Ab-mediated tethering. The light pink curve indicates the average plasma cell output 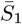, and the solid black curves correspond to contours of 5%, 50%, and 95% probability of Ag extraction. Parameters: *λ*_1_ = 2.9; *S*_dir_ = 1; *S*_unbound_ = 10^4^; *A*_indir_ = 0 (i.e., no Ab-mediated tethering).

However, the preceding situation considers only a single epitope: what if a B cell lineage targeting a second epitope is present? One might expect that, if affinity maturation relies on masking alone, then a second B cell lineage targeting a non-overlapping epitope could outcompete lineage 1 even if lineage 2 is weaker, because it does not face the obstacle of masking. Fig. 3a (left) shows a kymograph of a GC simulation with two non-overlapping epitopes, in which B cells targeting epitope 1 (red) have a notably stronger precursor distribution (*λ*_1_ = 2.5) than those targeting epitope 2 (blue; *λ*_2_ = 1.7). Initially, B cells targeting epitope 1 mature in strength, raising the 5%-to-95% window for lineage 1; however, the 5%-to-95% window for B cells targeting epitope 2 still has not moved, because as a non-overlapping epitope it is not affected by 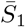. This allows lineage 2 to establish and then dominate the GC reaction, in this case driving lineage 1 to extinction.

**FIG. 3.**
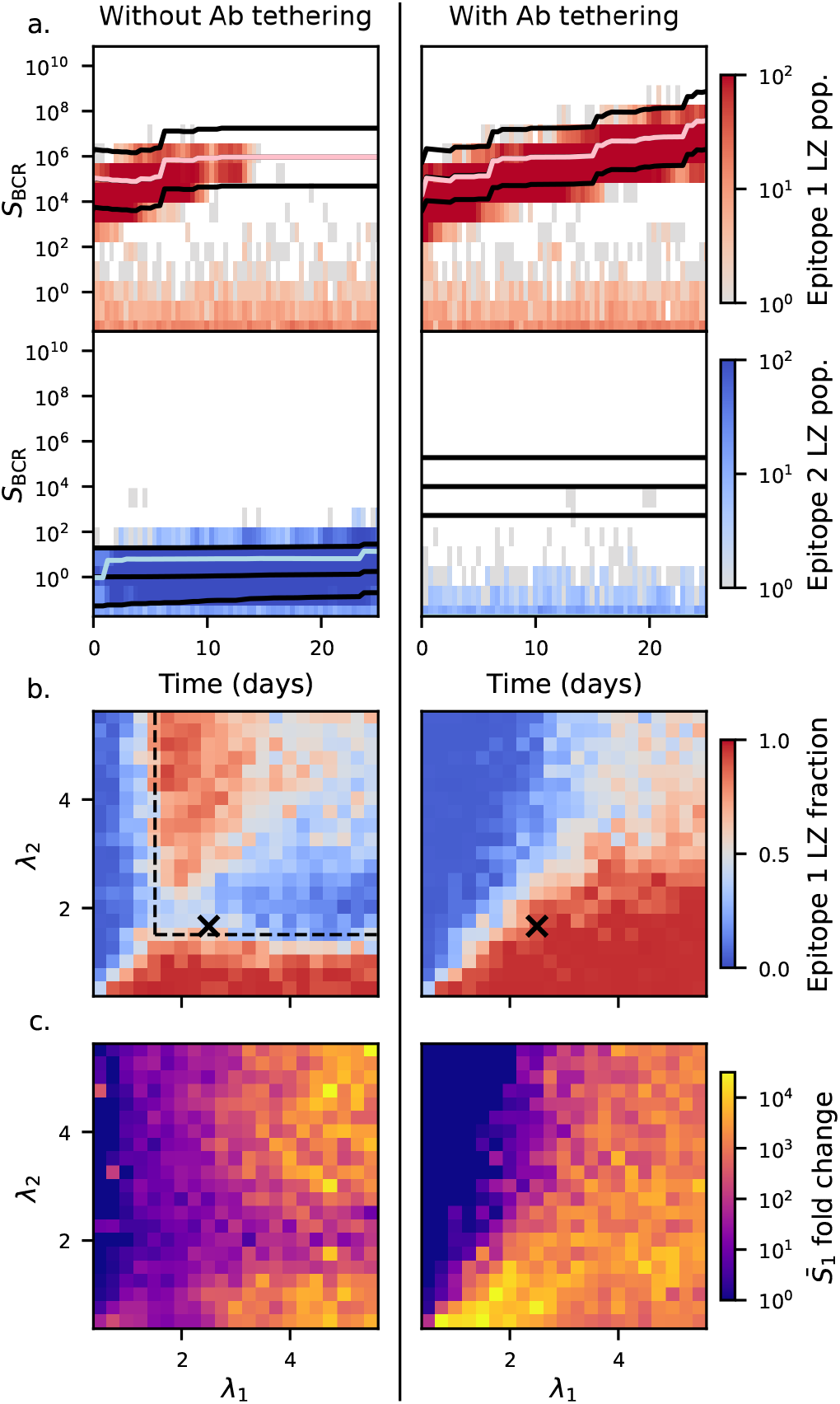
Ab-mediated tethering stabilizes affinity maturation in the presence of multiple epitopes. **(a)** Example kymographs from GC simulations with and without Ab-mediated tethering for two competing epitopes. Precursor distributions are set by *λ*_1_ = 2.5 and *λ*_2_ = 1.7. Black curves: 5%, 50%, and 95% extraction-probability. Light pink and blue curves: 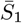 and 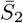, respectively. Without Ab-mediated tethering (left), low-strength B cells targeting epitope 2 (blue) outcompete resident higher-strength B cells targeting epitope 1 (red), which must compete against stronger epitope masking. With Ab-mediated tethering (right), epitope-1 B cells remain dominant and continue to increase in strength. **(b)** Heatmaps showing the fraction of LZ B cells targeting epitope 1 on day 25 as a function of the precursor-distribution decay scales (*λ*_1_, *λ*_2_). Without Ab-mediated tethering (left), inverted regions (red above the diagonal, blue below) emerge, where B cells targeting the epitope with weaker precursors outcompete those targeting the stronger precursors. Dashed black lines: theoretical estimate of the inversion boundary. With Ab-mediated tethering (right), the phase diagram collapses into two contiguous phases separated by the *λ*_1_ = *λ*_2_ diagonal, indicating sustained focus on the dominant precursor population. Aside from stochasticity, the diagram is symmetric about the diagonal. Each pixel represents the mean of 20 random initializations. The example in panel (a) is marked by *×*. **(c)** Fold change in mean Ab strength, 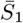, for epitope 1 between days 0 and 25. Ab-mediated tethering (right) yields more consistent gains in Ab strength. Parameters: *S*_dir_ = 1; *S*_unbound_ = 10^4^; *S*_indir_ = 0.2; *A*_indir_ = 0 (left), 10^4^ (right).

To examine the susceptibility of the GC to this kind of population inversion, Fig. 3b (left) shows a phase diagram parameterized by the decay scales of the precursor distributions (*λ*_1_, *λ*_2_) (recall, larger *λ*_*i*_ indicates stronger precursors), with color indicating the fraction of LZ B cells targeting epitope 1 on day 25.

Without Ab-mediated tethering, the phase diagram features three qualitatively distinct regions. When either *λ*_1_ or *λ*_2_ is sufficiently small, the GC exists in a *hierarchy-preserving* regime (blue above the diagonal, red below the diagonal) in which the B cells targeting the dominant epitope (larger *λ*_*i*_) dominate the GC. At slightly larger *λ*_1_ or *λ*_2_, the system enters an *inverted* regime (red above the diagonal, blue below the diagonal). In this regime, B cells targeting the subdominant epitope (smaller *λ*_*i*_) ultimately dominate instead. Finally, at sufficiently large (*λ*_1_, *λ*_2_), the GC enters an *equal-competing* regime (gray), in which B cells targeting either epitope are able to coexist. In Fig. 3c, we assess the extent of affinity maturation by comparing the fold increase in plasma-cell output, 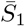, between day 0 and day 25. The inverted regime shows a notable reduction in plasma-cell strength fold change—potentially consequential as affinity maturation is most valuable when starting from modest precursor B cell strengths.

Mechanistically, a population inversion requires the precursors from the subdominant lineage to be sufficiently strong compared to *S*_dir_ such that they are not excluded from the GC reaction, but sufficiently weak compared to *S*_unbound_ that they do not immediately feel their Ab feedback. In the absence of Ab-mediated tethering, *S*_unbound_ is the only free parameter of the phenomenological Ab-feedback model; it sets the level of total plasma cell output, i.e., ∑_*α ∈*plasma(*i*)_ *S*_BCR,*α*_ in Eq. (2), at which masking begins to set in. In Supplementary Fig. S1, we explore the dependence of the phase diagram on *S*_unbound_: The upper limit of the inverted regime increases with *S*_unbound_, while the presence of an inverted regime requires *S*_unbound_ ≳ *S*_dir_ (cf. Fig. S1). We also note that the phase diagrams in Fig. 3 apply for well-separated epitopes; Supplementary Fig. S2 treats the case of epitope overlap, in which antibodies specific to one epitope can inhibit binding to a second epitope. The inverted regime persists under modest epitope overlap, but is naturally suppressed once Abs from the dominant epitope strongly block binding to the subdominant one.

Finally, in Fig. 3b (left), we include a theoretical estimate of the phase boundary between the inverted and hierarchy-preserving phases (dashed black lines). This estimate is based on a simplified picture in which the immunodominant epitope feels only masking and the sub-dominant epitope feels only direct tethering (see Supplementary Information). The agreement quantitatively corroborates these forces as the main mechanism driving the onset of the inverted regime in the absence of Ab-mediated tethering.

### Ab-mediated tethering broadens the region of continued affinity maturation

Continued affinity maturation on multi-epitope Ags requires a mechanism by which B cells targeting any epitope experience an increasing barrier to extracting Ag as the IC Ab strength for the immunodominant epitope increases. To this end, we now consider the full model with masking, direct tethering, and Ab-mediated tethering (*A*_indir_ *>* 0). Fig. 3a (right) shows a kymograph for the same parameters as the left panel, but now including Ab-mediated tethering (*S*_indir_ = 0.2 and *A*_indir_ = 10^4^). In notable contrast, lineage 2 no longer outcompetes lineage 1: Ab-mediated tethering shifts the 5%-to-95% window upwards for *both* epitope classes, effectively preventing the B cells targeting the subdominant epitope from successfully competing in the GC.

The avoidance of population inversion is evident in the phase diagram of Fig. 3b (right). In contrast to the left panel, the phase diagram now comprises just two contiguous regions separated by the diagonal—the B cell class with the higher precursor strength dominates the GC population. Beyond stabilizing GC composition in this way, we ask whether Ab-mediated tethering also improves the degree of affinity maturation: Fig. 3c shows that the fold increase of epitope 1 strength is greater and more consistent in the presence of Ab tethering. In the hierarchy-preserving regions (small *λ*_1_ and *λ*_2_), the increased affinity maturation is due to the fact that Ab-mediated tethering accelerates the rise of the 5%-to-95% window as a function of 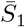 for epitope 1; in the previously inverted regions, the increased affinity maturation follows because the weaker lineage fails to invade the GC.

### Dynamic receptor expression may allow controlled GC diversification

The mechanism in Fig. 3 suggests that Ab-mediated tethering may focus GC selection on immunodominant epitopes. The strength of Ab-mediated tethering is likely determined in part by the availability of receptors that couple the IC-bound Abs to the FDCs. Indeed, one such receptor, Fc*γ*RIIb, which binds directly to the conserved tails of IgGs, is known to have dynamic expression on FDCs [50], raising the conceptual possibility that Ab-mediated tethering may be actively regulated. To explore this possibility, in Fig. 4a we consider a version of Eq. (3) in which the amplitude *A*_indir_(*t*) is time-dependent. In particular, we consider a simple scenario where *A*_indir_(*t*) is a step function from a maximum value before *t* = *t*_c_ to zero afterwards. Initially, the B cells targeting the subdominant epitope are blocked from participating, but once expression of the indirect tethering proteins is reduced, the effective barrier to entry for these B cells falls as well, allowing the GC to diversify. Fig. 4b compares the fraction of the GC targeting epitope 1 between days 20 and 50, showing that the dynamic reduction of *A*_indir_(*t*) is accompanied by the enlargement of the inverted and equal-competing regimes.

**FIG. 4.**
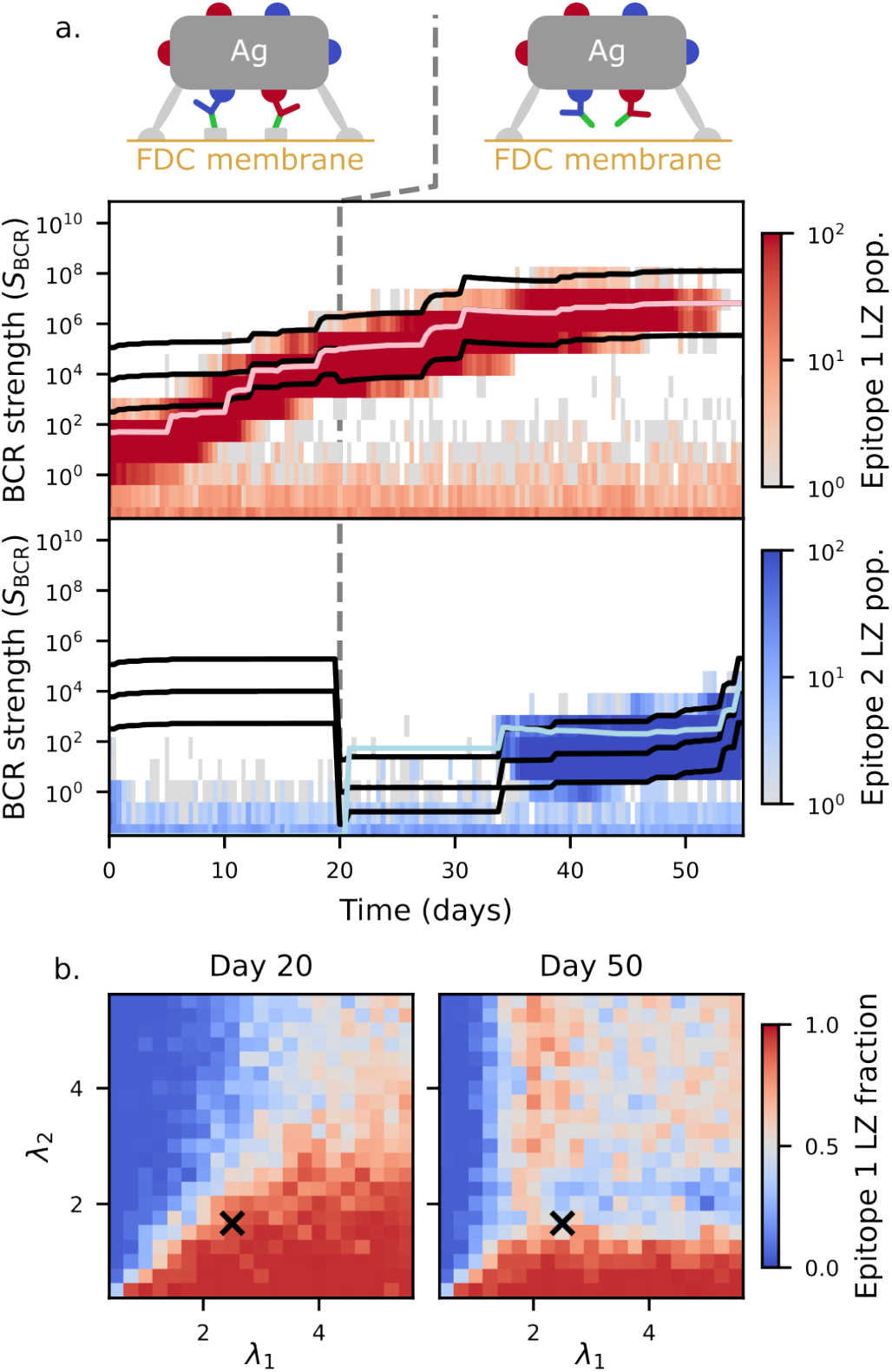
Steering GC diversity via Ab-tethering receptors. **(a)** Upper: schematic of downregulation of Ab-mediated tethering proteins on an FDC. Lower: kymograph of GC dynamics with time-dependent Ab tethering. In this example, the Ab-tethering magnitude is a step function *A*_indir_(*t*) = 10^4^Θ(*t*_c_ *™ t*) with *t*_c_ = 20 days, where Θ is the Heaviside step function. The decrease in Ab tethering lowers the effective barrier to entry for epitope-2 B cells, allowing diversification of the GC. **(b)** Heatmaps of the LZ composition on day 20 and day 50. Each pixel represents the mean of 20 random initializations. Parameters: *S*_dir_ = 1; *S*_indir_ = 0.2; *S*_unbound_ = 10^4^. The *×* marks location (*λ*_1_, *λ*_2_) = (2.5, 1.7) of the example in (a).

This situation is qualitatively similar to observed trends in longer GC reactions, which initially drive affinity maturation, but ultimately give rise to diversification [24, 40, 61]. A common interpretation of these experiments is that masking effects, potentially along with increased polymerization of ICs [24], naturally give rise to diversification in a largely unregulated fashion. Our results raise the possibility that FDCs could also actively shape the timing and extent of this diversification through receptor regulation. The extent to which this occurs remains to be explored experimentally.

## Discussion

Here we presented a biophysical picture in which two mechanisms of Ag tethering on FDCs collaborate to regulate the B cell epitope diversity generated by the GC reaction. An implication of this model is that the absence of dedicated Ab receptors on FDCs, such as Fc*γ*R, should result in GCs that produce relatively low affinity but highly diverse B cell repertoires. This is consistent with experiments [62] in which mice lacking Fc*γ*Rs on FDCs show reduced affinity but increased diversity, although the extent to which this diversity was attributable to epitope diversity specifically was not quantified. Interestingly, the magnitude of the effect in these experiments was Ag dependent. Because the proportion of direct versus Ab-mediated tethering (either through Fc*γ*Rs or Ab-bound complement) likely depends on availability of complement and Ab binding sites, additional investigation into the relative availability of these sites would be informative.

These Fc*γ*R knockout experiments could be complemented by additional, more nuanced probes. For instance, consider immunization with an Ag featuring two well-separated epitopes, *E*_1_ and *E*_2_. If IgG Abs with high affinity for *E*_1_ are administered before and during immunization, Ab-mediated tethering of Ag to FDCs by Fc*γ*R bonds will suppress the immune response for both *E*_1_ and *E*_2_. Suppose that instead of IgG, one administers F(ab)_2_, which lacks an Fc tail, and so is not susceptible to tethering via Fc*γ*Rs. In this case, our mechanism would imply that development of Abs for *E*_1_ may be suppressed due to masking by the external F(ab)_2_, but affinity maturation for *E*_2_ could still proceed. Versions of this experiment could also be done with other non-IgG immunoglobulins.

In addition to their mechanical roles in Ag extraction, other functional roles for CRs and Fc*γ*Rs on FDCs have been demonstrated or proposed. The expression level of CRs on FDCs is relatively constant over time, and these receptors have been implicated in defining the spatial organization of the LZ [63], retaining Ag for long periods of time [5], and receiving Ag from noncognate B cell escorts [64]. The precise roles of Fc*γ*R are a subject of investigation. Fc*γ*R is also expressed by GC B cells and has an inhibitory effect in that context. It has therefore been suggested that one role of Fc*γ*Rs on FDCs is to sequester Fc tails away from B cells, thereby making ICs on the surface of FDCs more immunopotent [50]. Moreover, Fc*γ*R expressed on FDCs appears to support FDC activation, including chemokine release and further upregulation of Fc*γ*R [51]. The role of Ag extraction considered here also needs to be disentangled from several other factors influencing B cell selection, including time-evolving follicular helper and regulatory T cell populations [65–67], changes in Ag levels [37, 68], and re-entrance of circulating memory B cells [40].

Finally, our model suggests the intriguing possibility that FDCs could dynamically regulate the affinity– diversity trade-off over time. While Ab feedback is to a large extent unavoidable, it is plausible that GCs have evolved ways to actively steer its influence. Unlike the CRs, whose expression remains roughly constant over time, Fc*γ*R is upregulated roughly 50-fold upon FDC activation [50]. This early upregulation may initially focus the GC reaction on nurturing one immunodominant epitope class. A later downregulation would then permit epitope diversification over longer time scales. A crossover from affinity increase early in the GC reaction to increased diversity later is indeed generally observed in longer GC experiments [24, 40, 61]; however, direct connection to temporal modulation of Fc*γ*R expression on FDCs has not, to our knowledge, been systematically studied. We hope our work inspires further investigation into the biophysical mechanisms regulating affinity and diversity in the GC reaction.

## Acknowledgments

The authors would like to thank MC Carroll, N Claussen, J Merkenschlager, MC Nussenzweig, and J Yuly for valuable discussions. AGT Pyo was supported by the Natural Sciences and Engineering Research Council of Canada and a Stanford Science Fellowship. DS was supported by the Human Immunome Project and Michelson Medical Research Foundation’s Michelson Prize: Next Generation Grant. This work was supported in part by Princeton University through the Center for the Physics of Biological Function. This work was also supported by the Weill Cancer Hub East through funding to NSW. This work was performed in part at Aspen Center for Physics, which is supported by National Science Foundation grant PHY-2210452.

## Methods

Here we provide the details of our minimal model for the GC reaction. The agent-based model consists of a population of GC B cells, each of which at any given time is in one of three compartments: light zone (LZ), dark zone (DZ), or plasma B cell compartment. Each B cell is also assigned an epitope class that it targets, and its B cell receptor (BCR) has a phenomenological strength *S*_BCR_ that quantifies its ability to extract Ag from FDCs. Note that we opt to use an effective parameter *S*_BCR_, rather than a specific physical parameter, such as equilibrium dissociation constant or binding time, because Ag extraction is a complex nonequilibrium process involving force, mechanics, and complicated geometries. Nonetheless, one may think of *S*_BCR_ as being roughly related to BCR affinity.

In each time step d*t*, the following processes occur:

1. **Apoptosis:** Each LZ B cell has a probability *r*_*ϕ*_ d*t* of undergoing apoptosis.
2. **Ag acquisition:** Each LZ B cell has a probability *r*_FDC_ d*t* of interacting with an FDC. Upon interaction, a B cell with strength *S*_BCR_ for epitope *i* extracts one unit of Ag with probability *p*(*S*_BCR_, *i*) prescribed by the phenomenological description in Eqs. (1-4). For simplicity, we assume a constant supply of Ag.
3. **Selection by T cells:** To impose positive selection signals on LZ B cells, we use a simple model of T cell help. In the time step d*t*, each LZ B cell has a probability 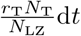 of interacting with a T cell, where *N*_LZ_ is the total number of B cells in the LZ and *N*_T_ is the total number of T cells, taken to be constant. Upon interaction, a B cell carrying *a* units of Ag is assigned *D*(*a*) divisions. If *D*(*a*) = 0, then the B cell continues in the LZ. If *D*(*a*) *>* 0, then the B cell has a probability *p*_ex_ of exiting the GC as a single plasma B cell and a probability (1 ™ *p*_ex_) of entering the DZ for divisions. The functional form of *D*(*a*) is

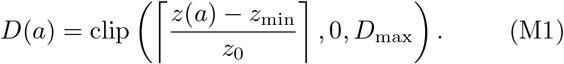

Here, *z*(*a*) is the *z*-score associated with the Ag levels in the LZ

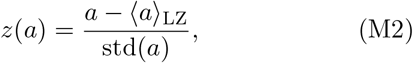

where *a* _LZ_ and std(*a*) are the mean and standard deviation, respectively, of Ag per B cell over the LZ at time *t*. In Eq. (M1), *z*_min_ sets the *z*-score at which nonzero divisions are assigned, 1*/z*_0_ sets the slope, and *D*_max_ is the maximum number of divisions allowed.
4. **Division and mutation:** In the DZ, each B cell is labeled by its number of remaining divisions, initially *D*(*a*). Each B cell has a probability *r*_div_d*t* of dividing in each time step. When the number of remaining divisions reaches 0, each daughter acquires a single mutation with probability *p*_mut_. The mutation increases or decreases the log strength *s* = log *S*_BCR_ by step size *δs* with probability *p*_+_ or (1 ™ *p*_+_), respectively. The B cell then returns to the LZ, where it starts with *a* = 0 units of Ag.
5. **Entrance of naïve B cells:** With probability *r*_ent_d*t* a new B cell enters the LZ. The B cell targets epitope *i* with probability *p*_*i*_. For simplicity, we take *p*_*i*_ = 1*/N*_ep_ where *N*_ep_ is the number of epitopes. We assign the log strength *s* = log *S*_BCR_ of the entering B cell by sampling from a distribution *ρ*_*i*_(*s*) that depends on the epitope *i*. We take *ρ*_*i*_(*s*) to be a discrete exponential distribution [44]:

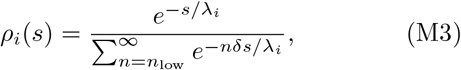

with support on the semi-infinite lattice

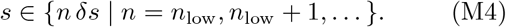

In practice, we take the lower cutoff to be *n*_low_ = ™ 3, which is sufficiently low (given the parameters described below) that any B cell with an log strength lower than *δs n*_low_ is effectively unable to acquire Ag and is therefore naturally excluded from the GC.

To initialize the simulation, we draw *N*_0_ = 100 B cells from the precursor distribution, i.e., Eq. (M3), with an equal number targeting each epitope. The initial B cells start in the LZ, and we define initialization as day ™ 5. We allow the system to equilibrate for 5 days, which is enough time for the GC population to stabilize and to allow selection to act on variance arising from mutations rather than solely from the precursor distribution.

The parameters used in the simulations are summarized in Methods Table M1, and are chosen to exhibit stable GC dynamics with a steady-state population between 1,000 and 3,000 B cells. The values of the parameters for the masking and tethering, namely *S*_unbound_, *S*_dir_, *S*_indir_, and *A*_indir_ appearing in Eqs. (1-3), are listed in the relevant figure captions. Without loss of generality we set *S*_dir_ = 1. For simplicity, we take *S*_indir_ = 0.2 *< S*_dir_, which implies that the factor 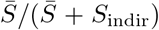 in Eq. (3) is quickly saturated, typically by day 1 of the GC simulation. Variation in *S*_unbound_ and *A*_indir_ as well as the effects of epitope overlap are explored in the Supplementary Information.

**TABLE M1.** Model parameters. Values for *S*_unbound_, *S*_dir_, *S*_indir_, *A*_indir_ vary across simulations and are listed in the relevant captions. Without loss of generality, we take *S*_dir_ = 1 to set an overall scale.

| Parameter | Description | Value |
| --- | --- | --- |
| $r_\phi$ | Apoptosis rate | $(1/6) \text{ hr}^{-1}$ |
| $r_{\text{FDC}}$ | FDC interaction rate | $20 \text{ hr}^{-1}$ |
| $N_{\text{T}} r_{\text{T}}$ | Maximum T cell help rate | $10 \text{ hr}^{-1}$ |
| $z_{\text{min}}$ | Division assignment Eq. (M1), $z$-threshold | 0.0 |
| $z_0$ | Division assignment Eq. (M1), scale | 0.3 |
| $D_{\text{max}}$ | Division assignment Eq. (M1), maximum divisions | 9 |
| $r_{\text{div}}$ | Division rate | $(1/5) \text{ hr}^{-1}$ |
| $p_{\text{ex}}$ | Export probability | 0.05 |
| $r_{\text{ent}}$ | Entrance rate | $20 \text{ hr}^{-1}$ |
| $p_i$ | Entrance probability per epitope | $1/N_{\text{ep}} = 1/2$ |
| $p_{\text{mut}}$ | Mutation probability | 0.6 |
| $p_+$ | Beneficial mutation fraction | 0.01 |
| $\delta s$ | Logarithmic mutation step size | 2.0 |

