## Supplementary Information for "Antibody-mediated antigen tethering as a regulator of germinal-center B cell affinity and diversity"


### FIRST-PASSAGE MODEL FOR ANTIGEN EXTRACTION

In this section, we present a toy microscopic model of Ag extraction based on force-free first-passage-time dissociation. We emphasize that this first-passage model is intentionally simple, ignoring important aspects such as molecule geometry, flexibility, membrane mechanics, and pulling force. It nonetheless exemplifies the key scales appearing in the phenomenologically motivated $p(S_{\rm BCR}, i)$, Eq. (1), in the main text. As a simple first case to illustrate direct tethering, we consider a single antigen (Ag) molecule bound to the FDC membrane by a single direct bond (e.g., via complement bound to a complement receptor (CR)), and to the B cell membrane by a single B cell receptor (BCR) bond. In this force-free first-passage picture, we say that the B cell "extracts" the Ag if the CR bond breaks before the BCR bond breaks. Assuming that the detachment times are exponentially distributed with bond lifetimes $\tau_{\rm CR}$ and $\tau_{\rm BCR}$, the probability of extraction is

$$P_{\rm extract} = \frac{\text{Rate of FDC bond breaking}}{\text{Rate of either bond breaking}} \tag{S1}$$

$$= \frac{1/\tau_{\rm CR}}{(1/\tau_{\rm CR}) + (1/\tau_{\rm BCR})} \tag{S2}$$

$$= \frac{1}{1 + \tau_{\rm CR}/\tau_{\rm BCR}}. \tag{S3}$$

Notice that Eq. (S3) has the form of a direct-tethering sigmoid if one identifies $\tau_{\rm BCR}$ with $S_{\rm BCR}$ and $\tau_{\rm CR}$ with $S_{\rm dir}$. To include masking, now suppose that the BCR must compete with a soluble IgG Ab to bind to the Ag. A simple formula for the extraction probability is the probability that the BCR rather than the IgG is bound to the Ag times the probability of extraction if the Ag is indeed bound by the BCR. Letting $\tau_{\rm IgG}$ denote the bond lifetime for the IgG-Ag bond, we can write

$$P_{\rm extract} = (\text{Probability BCR bound to Ag}) \times (\text{Probability of extraction if BCR bound to Ag}) \tag{S4}$$

$$= \underbrace{\left(\frac{1}{1 + \tau_{\rm IgG}/\tau_{\rm BCR}}\right)}_{\text{Masking}} \underbrace{\left(\frac{1}{1 + \tau_{\rm CR}/\tau_{\rm BCR}}\right)}_{\text{Direct tethering}}. \tag{S5}$$

The first term in Eq. (S5) can be physically interpreted as the fraction of the amount of time the Ag spends bound to the BCR vs. bound to the IgG in a simple equilibrium binding scenario with equal attachment rates, and negligible time spent unbound to both. Notice that Eq. (S5) has the same functional form as our phenomenological model with Ab masking and direct tethering if one identifies $\tau_{\rm IgG}$ with $\bar{S}_i$, $\tau_{\rm BCR}$ with $S_{\rm BCR}$, and $\tau_{\rm CR}$ with $S_{\rm dir}$.

As a generalization, now suppose that the Ag immune complex (IC) may be bound to the membrane of each cell by multiple bonds, so we must account for the fact that individual bonds can rebind. To address this, first consider a generic system of $N$ bonds with bond lifetimes $\tau_i$ and rebinding times $\tau_{{\rm off},i}$, where $i = 1 \ldots N$. Our interest is in the limit in which rebinding is much faster than unbinding, i.e. $\tau_{{\rm off},i}/\tau_j \ll 1$ for all $i, j$. Then the collective off-time $\tau_{\rm off,total}$, i.e. the average time it takes the first bond to form starting from the completely unbound state, is

$$\frac{1}{\tau_{\rm off,total}} = \sum_i \frac{1}{\tau_{{\rm off},i}}. \tag{S6}$$

The total on-time $\tau_{\text{total}}$, i.e., the average time to go from a bound state to the completely unbound state, is obtained via the relationship

$$\frac{\tau_{\text{off,total}}}{\tau_{\text{total}}} \approx \prod_i \frac{\tau_{\text{off},i}}{\tau_i}. \tag{S7}$$

Equation (S7), valid in the fast rebinding limit, can be obtained by noting that the left-hand side represents the fraction of time that no bond is bound, and the right-hand side decomposes this fraction into the product over individual bonds.

For simplicity, in what follows, we will assume that all bonds rebind with approximately similar rebinding times, denoted simply $\tau_{\text{off}}$. Suppose the B cell membrane is tethered to the IC by $N_{\text{BCR}}$ BCR bonds. From Eqs. (S6-S7), the time constant $\tau_{\text{B cell}}$ for detachment of the B cell is given by

$$\frac{1}{\tau_{\text{B cell}}} = \frac{N_{\text{BCR}}}{\tau_{\text{off}}} \left(\frac{\tau_{\text{off}}}{\tau_{\text{BCR}}}\right)^{N_{\text{BCR}}}. \tag{S8}$$

On the FDC, we assume that the direct tethering is provided by $N_{\text{CR}}$ CRs, and that the Ab-mediated tethering is provided by $N_{\text{Fc}\gamma\text{R}}$ composite Fc$\gamma$R-IgG-Ag bonds. In this case, the time constant $\tau_{\text{FDC}}$ is given by

$$\frac{1}{\tau_{\text{FDC}}} = \frac{N_{\text{CR}} + N_{\text{Fc}\gamma\text{R}}}{\tau_{\text{off}}} \left(\frac{\tau_{\text{off}}}{\tau_{\text{CR}}}\right)^{N_{\text{CR}}} \left(\frac{\tau_{\text{off}}}{\tau_{\text{IgG}}} + \frac{\tau_{\text{off}}}{\tau_{\text{Fc}\gamma\text{R}}}\right)^{N_{\text{Fc}\gamma\text{R}}}, \tag{S9}$$

where $\tau_{\text{Fc}\gamma\text{R}}$ is the time constant associated with the IgG-Fc$\gamma$R bond. We have used the fact that the detachment rate of the composite Fc$\gamma$R-IgG-Ag bond is

$$\frac{1}{\tau_{\text{composite}}} = \frac{1}{\tau_{\text{IgG}}} + \frac{1}{\tau_{\text{Fc}\gamma\text{R}}}. \tag{S10}$$

The probability that the FDC detaches first is then given by

$$P_{\text{extract}} = \frac{1/\tau_{\text{FDC}}}{(1/\tau_{\text{FDC}}) + (1/\tau_{\text{B cell}})} \tag{S11}$$

$$= \underbrace{\frac{1}{1 + (\tau_*/\tau_{\text{BCR}})^{N_{\text{BCR}}}}}_{\text{Tethering}}, \tag{S12}$$

where

$$\tau_* = \tau_{\text{off}} \left(\frac{N_{\text{BCR}}}{N_{\text{CR}} + N_{\text{Fc}\gamma\text{R}}}\right)^{\frac{1}{N_{\text{BCR}}}} \left(\frac{\tau_{\text{CR}}}{\tau_{\text{off}}}\right)^{\frac{N_{\text{CR}}}{N_{\text{BCR}}}} \left(\frac{\tau_{\text{IgG}}/\tau_{\text{off}}}{1 + \tau_{\text{IgG}}/\tau_{\text{Fc}\gamma\text{R}}}\right)^{\frac{N_{\text{Fc}\gamma\text{R}}}{N_{\text{BCR}}}}. \tag{S13}$$

Note that Eq. (S13) is derived under the assumption that $\tau_{\text{off}}/\tau_{\text{IgG}} \ll 1$. In the opposite limit, the Ag is effectively only tethered to the FDC by the CR bonds, so one obtains

$$\tau_* = \tau_{\text{off}} \left(\frac{N_{\text{BCR}}}{N_{\text{CR}}}\right)^{\frac{1}{N_{\text{BCR}}}} \left(\frac{\tau_{\text{CR}}}{\tau_{\text{off}}}\right)^{\frac{N_{\text{CR}}}{N_{\text{BCR}}}}. \tag{S14}$$

For a simple expression that interpolates between both limits, we first neglect the prefactors that depend only on $N_{\text{BCR}}$, $N_{\text{CR}}$, and $N_{\text{Fc}\gamma\text{R}}$ because they are negligible on a logarithmic scale and ultimately independent of the time scales of interest, $\tau_{\text{IgG}}$, $\tau_{\text{off}}$, $\tau_{\text{Fc}\gamma\text{R}}$. Second, we regularize the numerator of the final factor in Eq. (S13) such that the factor approaches 1 as $\tau_{\text{IgG}} \to 0$. Using these two approximations, we obtain

$$\tau_* \approx \tau_{\text{off}} \left(\frac{\tau_{\text{CR}}}{\tau_{\text{off}}}\right)^{\frac{N_{\text{CR}}}{N_{\text{BCR}}}} \left(\frac{1 + \tau_{\text{IgG}}/\tau_{\text{off}}}{1 + \tau_{\text{IgG}}/\tau_{\text{Fc}\gamma\text{R}}}\right)^{\frac{N_{\text{Fc}\gamma\text{R}}}{N_{\text{BCR}}}}. \tag{S15}$$

Notice that if one takes $N_{\text{CR}} = N_{\text{BCR}} = N_{\text{Fc}\gamma\text{R}} = 1$, one obtains the same functional form as Eq. (3) in the main text if one identifies $S_{\text{dir}}$ with $\tau_{\text{CR}}$; $\bar{S}$ with $\tau_{\text{IgG}}$; $S_{\text{indir}}$ with $\tau_{\text{Fc}\gamma\text{R}}$; and $A_{\text{indir}}$ with $(\tau_{\text{Fc}\gamma\text{R}}/\tau_{\text{off}}) - 1$. A masking prefactor can be included in Eq. (S12) as in Eq. (S5) to obtain the full functional form in Eq. (1) of the main text.

### MODEL DEPENDENCE ON FEEDBACK PARAMETERS

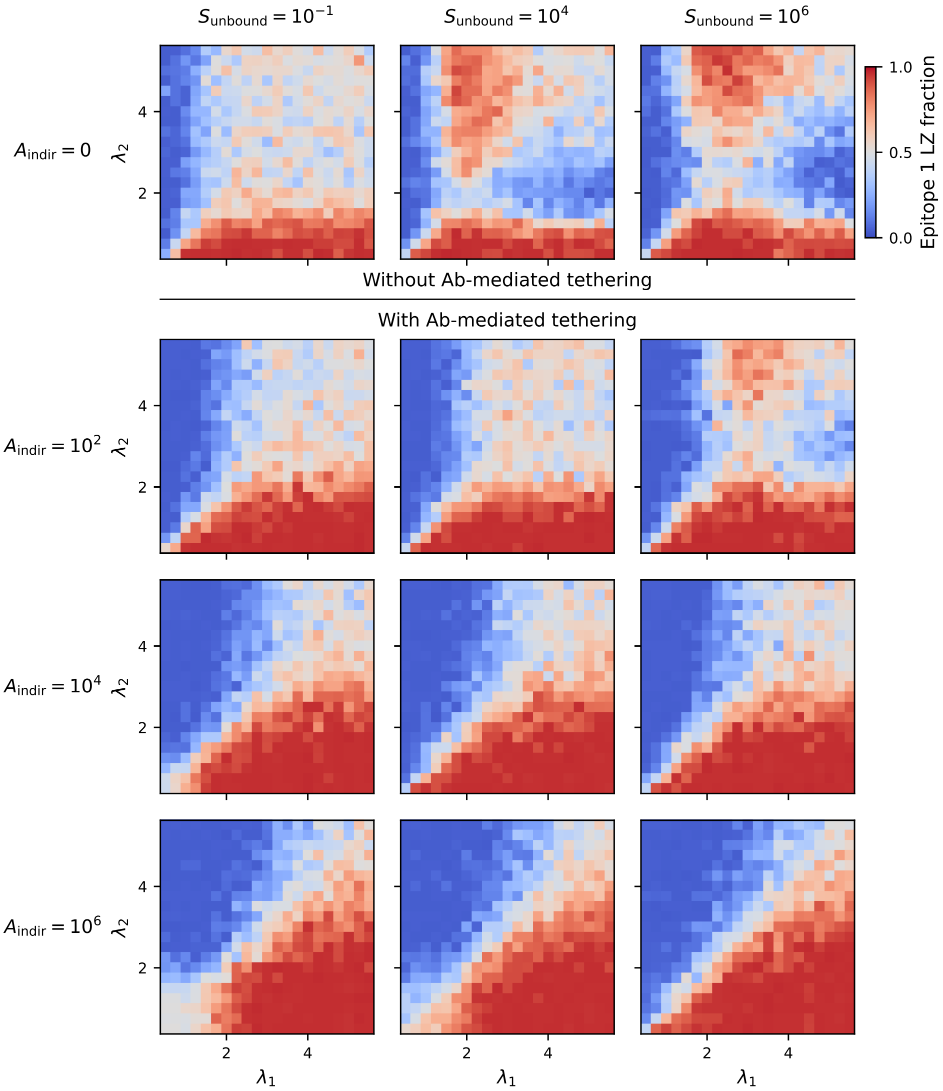


FIG. S1. **Dependence of GC composition on key model parameters.** Heatmaps showing the fraction of LZ B cells targeting epitope 1 on day 25 as a function of the precursor-distribution decay scales $(\lambda_1, \lambda_2)$, with variation over two key parameters of the model $S_{\mathrm{unbound}}$ and $A_{\mathrm{indir}}$. Recall that $S_{\mathrm{unbound}}$, appearing in Eq. (2), sets the strength at which the total plasma output, $\sum_{\alpha \in \mathrm{plasma}(i)} S_{\mathrm{BCR},\alpha}$, begins to meaningfully contribute to masking. The parameter $A_{\mathrm{indir}}$, appearing in Eq. (3), sets the maximum shift in tethering scale $S_{\mathrm{tether}}$ due to Ab-mediated tethering. Within each heatmap, there are up to three regimes: (1) a hierarchy-preserving regime (blue above the diagonal, red below the diagonal) in which the epitope with larger $\lambda_i$ (i.e. stronger precursors) dominates the GC; (2) an inverted regime (red above the diagonal, blue below the diagonal) in which the epitope with smaller $\lambda_i$ (i.e. weaker precursors) dominates the GC; and (3) an equal-competing regime (gray) in which the epitopes coexist. Without Ab-mediated tethering ($A_{\mathrm{indir}} = 0$), the existence of an inverted regime requires $S_{\mathrm{unbound}} \gtrsim S_{\mathrm{dir}}$ so that masking is initially weak when precursor strengths are below $S_{\mathrm{dir}}$. Increasing $S_{\mathrm{unbound}}$ increases the size of the inverted regime. When $S_{\mathrm{unbound}} \not\gtrsim S_{\mathrm{dir}}$, the hierarchy-preserving and equal-competing regimes share a boundary. Depending on $S_{\mathrm{unbound}}$, a sufficiently large $A_{\mathrm{indir}}$ is required in order to suppress the inverted regime. Increasing $A_{\mathrm{indir}}$ also decreases the equal-competing regime. In all plots, $S_{\mathrm{indir}} = 0.2$ and $S_{\mathrm{dir}} = 1$; see Methods for discussion and Methods Table M1 for other model parameters.

### SENSITIVITY TO EPITOPE OVERLAP

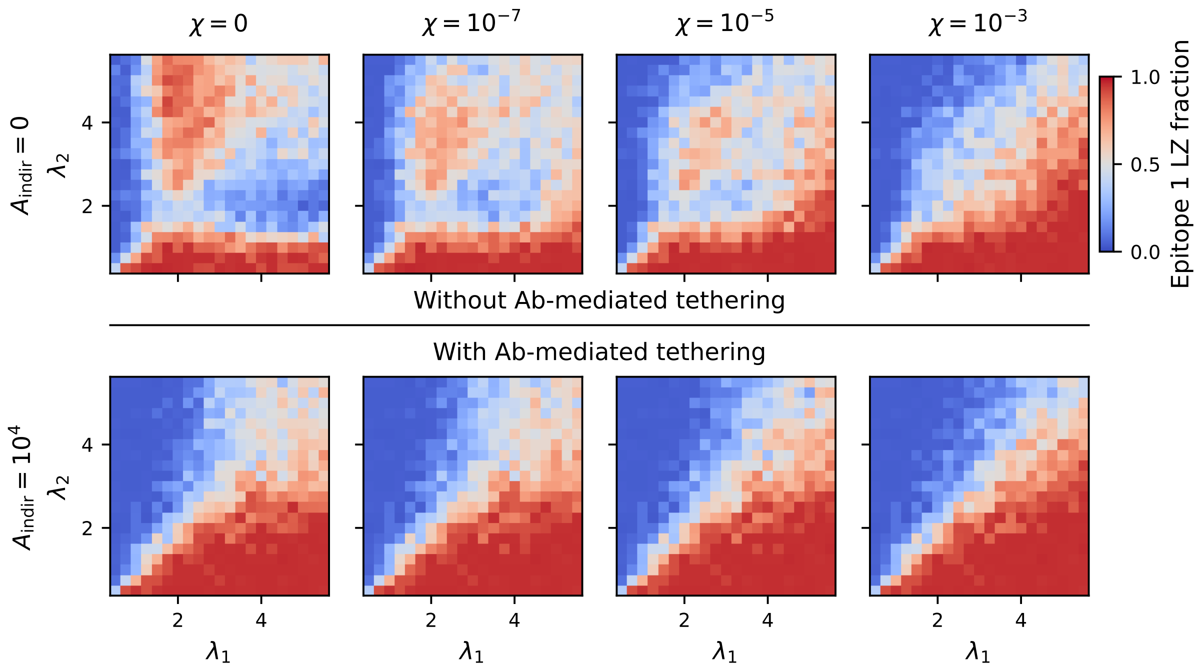


FIG. S2. **GC reaction with epitope overlap.** Heatmaps showing the fraction of LZ B cells targeting epitope 1 on day 25 as a function of the precursor-distribution decay scales $(\lambda_1, \lambda_2)$, with variation over the degree of epitope overlap, $\chi$. Top row: without Ab-mediated tethering; bottom row: with Ab-mediated tethering. See supplemental text for discussion. Parameters: $S_{\text{dir}} = 1$; $S_{\text{unbound}} = 10^4$; $S_{\text{indir}} = 0.2$.

In the main text, we focus on well-separated epitopes in which Abs bound to one epitope do not interfere with BCR binding to another epitope. Here we address the case of epitope overlap, in which partial occlusion occurs. Phenomenologically, the probability of Ag extraction now takes the form

$$p(S_{\text{BCR}}, i) = \sigma(S_{\text{BCR}}/S_{\text{tether}})\sigma(S_{\text{BCR}}/M_i), \tag{S16}$$

where the masking scale $M_i = \sum_j \chi_{ij}\bar{S}_j$ is now given by a linear combination of the plasma output scales $\bar{S}_j$. We consider a simple parameterization of the coefficients in which self-masking is set to unity, $\chi_{11} = \chi_{22} = 1$, and the degree of epitope overlap is set by the off-diagonal element $\chi_{12} = \chi_{21} = \chi$. In Fig. S2, we vary $\chi$ by orders of magnitude, and examine its effects on GC outcomes. For modest epitope overlap, the inverted regime persists in the absence of Ab-mediated tethering, and epitope overlap acts to limit its extent at large $\lambda_i$. For stronger epitope overlap, the inverted regime is suppressed as cross-epitope masking is sufficient to suppress B cells targeting subdominant epitopes. Ab-mediated tethering continues to suppress the inverted regime.

### THEORETICAL ESTIMATE OF THE INVERSION PHASE BOUNDARY

Here we provide the details of the theoretical estimate of the phase boundary separating the hierarchy-preserving regime from the inverted regime in Fig. 3b (left).

The first step of the calculation is to perform simplified simulations in which only minimal ingredients are included. We initialize a simulation of the GC reaction in which only B cells targeting the immunodominant epitope (call it epitope 1 for concreteness) are present. We impose that B cells targeting this epitope compete only with masking effects, i.e. the probability of extracting Ag is given by

$$p(S_{\rm BCR}, 1) = \sigma(S_{\rm BCR}/\bar{S}_1), \tag{S17}$$

where $\bar{S}_1$ is given by Eq. (2) in the main text. We take $S_{\rm unbound} = 0$ for these simulations, so that the B cells targeting epitope 1 undergo affinity maturation in a landscape that is translationally invariant in $\log S_{\rm BCR}$. We allow the simulation to run for a sufficient amount of time for the population to reach a hill-climbing steady state, and no external B cells targeting any epitope are introduced during this period.

Deep in the steady state of the simulation (30 days), we instantaneously inject into the LZ $M$ identical B cells targeting a subdominant epitope (epitope 2), each with strength $S_{\rm BCR}$. Unlike the B cells targeting epitope 1, the invading B cells are assumed to compete only with direct tethering, unaffected by masking: their probability of extracting Ag is given by

$$p(S_{\rm BCR}, 2) = \sigma(S_{\rm BCR}/S_{\rm dir}). \tag{S18}$$

We then allow the simulation to proceed with no further external B cells introduced, and check whether B cells targeting epitope 2 become established, defined in practice as not being extinct after 30 days. We repeat this for different values of injected strength $S_{\rm BCR}$, each with $N$ stochastic replicates, and we let $X$ denote the number of replicates in which B cells targeting the second epitope successfully establish. Approximating the $M$ injected B cells as independent, we estimate the probability $\mu(s)$ that a single B cell injected with log strength $s = \log S_{\rm BCR}$ for the epitope establishes:

$$\mu(s) = 1 - \left(1 - \frac{X}{N}\right)^{1/M}. \tag{S19}$$

The measured function $\mu(s)$ is shown in Supplementary Fig. S3a, with $N = 500$ and $M = 100$.

Taking $\mu(s)$ from the simplified simulations as an input, we now compute the prediction for the phase boundary $\lambda_{\rm crit}$ separating the hierarchy-preserving and inverted regimes in the full simulations. Recall that in the full simulations, naïve B cells targeting epitope $i$ are injected into the GC at a rate $p_i r_{\rm ent}$ and have a log strength $s = \log S_{\rm BCR}$ drawn from the distribution $\rho_i(s; \lambda_i)$, where larger strength-decay scale $\lambda_i$ indicates a higher average strength (see Methods). Assume for concreteness that $\lambda_1 > \lambda_2$, so that epitope 1 is thought of as the immunodominant epitope. Approximating the entering B cells as independent, one predicts that the probability that at least one B cell targeting epitope 2 establishes is

$$P_{\rm invert}(\lambda_2) = 1 - \exp\left[-N_2 \sum_s \rho_2(s; \lambda_2)\mu(s)\right], \tag{S20}$$

where $N_2 = p_2 r_{\rm ent} T_{\rm final}$ is the expected total number of injected B cells specific for epitope 2, and $T_{\rm final}$ is the length of the simulation. We define the phase boundary $\lambda_{\rm crit}$ as the solution to $P_{\rm invert}(\lambda_{\rm crit}) = \frac{1}{2}$. The curve $P_{\rm invert}(\lambda_2)$ is shown in Supplementary Fig. S3b for different values of $N_2$, adjusted via $T_{\rm final}$. Because $P_{\rm invert}$ is sufficiently sharp, the value of $\lambda_{\rm crit}$ depends only weakly on $N_2$. To compare against the full phase diagram, we take $p_2 = 1/2$, $T_{\rm final} = 25$ days, and $r_{\rm ent} = 20\,{\rm hr}^{-1}$. The resulting prediction ($\lambda_{\rm crit} = 1.51$) is shown in Fig. 3b and agrees well with the phase diagram based on the full simulations.

We emphasize that this calculation neglects several features of the full model. For example, in the simplified calculation, masking of the subdominant epitope is assumed to vanish exactly, although in the full model it will be nonzero (though small) once a single B cell is exported to the plasma fate. Invading B cells are treated as independent despite competing for limited T cell help. Finally, the initial conditions in the simplified model contain only B cells targeting the dominant epitope, whereas the full simulations begin with B cells targeting both epitopes.

The good agreement between theory and full simulation in light of these simplifications helps support the proposed inversion mechanism: population inversion occurs when a newly arriving B cell targeting the subdominant epitope

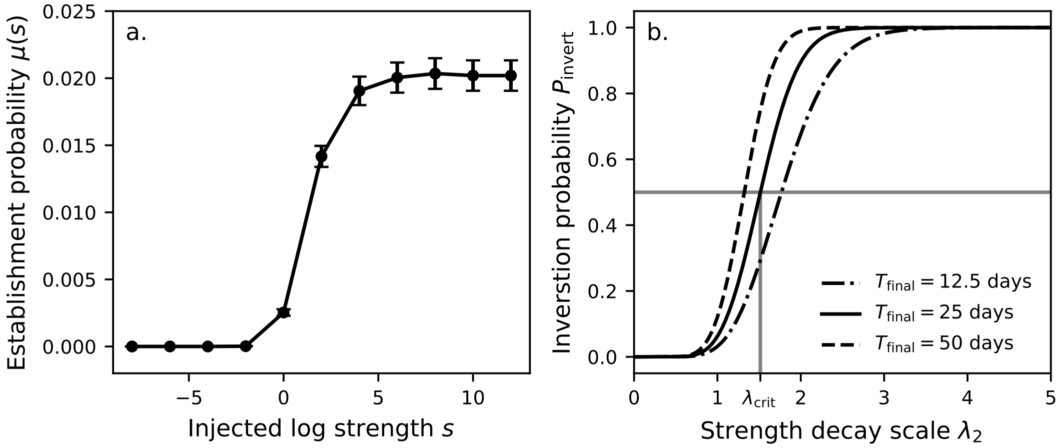


FIG. S3. **Theoretical derivation of phase boundary. (a)** The function $\mu(s)$ in Eq. (S19) measured with $M = 100$ injected B cells and $N = 500$ stochastic replicates. Error bars represent 1 standard deviation. **(b)** The function $P_{\mathrm{invert}}(\lambda_2)$ from Eq. (S20) with $p_2 = 1/2$, $r_{\mathrm{ent}} = 20\,\mathrm{hr}^{-1}$, and $T_{\mathrm{final}} = 25\,\mathrm{days}$ (solid line). Additional values of $T_{\mathrm{final}}$ illustrate relatively weak dependence of $\lambda_{\mathrm{crit}}$ on $T_{\mathrm{final}}$.

possesses a sufficiently large $S_{\mathrm{BCR}}$ relative to $S_{\mathrm{dir}}$, compared with the strength of the leading immunodominant clone relative to its masking scale $\bar{S}_1$. Moreover, the agreement in Eq. (S20) helps clarify how the phase diagrams of Fig. 3b depend on the parameters ($p_2$, $r_{\mathrm{ent}}$, and $\rho_i(s)$) related to the precursor distribution and entry rate.

*
†
